# Infection and transmission dynamics of bovine and human influenza A H5N1 viruses in mouse and hamster models

**DOI:** 10.64898/2026.01.17.700113

**Authors:** Mohammed Nooruzzaman, David W. Buchholz, Ruchi Rani, Pablo Sebastian Britto de Oliveira, Richard A. Adeleke, Chen Feng, Elena A. Demeter, Hector C. Aguilar, Diego G. Diel

## Abstract

Here we investigated the pathogenesis and contact transmission of bovine- and human-derived highly pathogenic avian influenza (HPAI) H5N1 clade 2.3.4.4b genotype B3.13 viruses in mammalian models. Using reverse genetics, we rescued three naturally occurring viruses: rTX2/24 (bovine-derived), rTexas/37 and rMichigan/90 (both human-derived), and compared their infection dynamics, replication and pathogenicity with the wild-type bovine TX2/24 strain *in vitro* and *in vivo*. All four viruses demonstrated comparable replication kinetics in four mammalian cell lines. However, the rMichigan/90 strain exhibited significantly smaller plaques in bovine and human cells. *In vivo* studies showed that mice infected with any of the viruses succumbed to infection within 4-5 days; however, mice infected with the rMichigan/90 virus exhibited slightly lower viral replication and shedding compared to the other strains. Similarly, as in the mouse experiments, in hamsters, all viruses induced body weight loss and oral shedding, with robust virus replication observed in tissues, but the rMichigan/90 virus presented reduced replication and shedding. Contact transmission studies in hamsters revealed limited transmissibility for these viruses, with only one out of four animals inoculated with the rMichigan/90 virus transmitting it to a naïve contact. These findings indicate that both bovine- and human-derived H5N1 genotype B3.13 viruses present high pathogenicity in mammals, though the overall transmissibility remains low.

**Significance Statement:** Influenza A H5N1 virus spilled over from wild birds into dairy cattle in the US in 2024. Since then, an increased number of human infections – with at least 71 confirmed cases – were confirmed. In the present study we evaluated and compared the pathogenicity and transmissibility of a bovine and two human H5N1 viruses of the genotype B3.13. Our results show that while all viruses are pathogenic in mouse and hamster models, the bovine isolate TX2/24 presents a broader tissue tropism than the human derived viruses. Contact transmission studies revealed a limited ability of these viruses to transmit in the hamster contact transmission model. These findings highlight the high pathogenicity of H5N1 viruses in mammals and demonstrate that that their transmissibility potential remains low.

## Introduction

The highly pathogenic avian influenza (HPAI) H5N1 virus clade 2.3.4.4b has become the globally dominant H5 lineage since 2021, causing an increasing number of mammalian spillover events across multiple continents. This clade has been linked to severe disease and high mortality in various mammalian species, particularly wild carnivores (1–4), aquatic mammals (5), and domestic cats (6–8). Histopathologic findings in mammals infected with this virus clade include severe central nervous system (CNS) inflammation with neuronal necrosis, pneumonia, and systemic inflammation (9). In early 2024, a reassortant genotype B3.13 virus of the clade 2.3.4.4b spilled over into dairy cows in Texas, United States (6, 10, 11) and subsequently spread rapidly in dairy cattle across the country (12), affecting more than 1,080 dairy herds in 18 states, as of December 13, 2025 (13). This epizootic with unprecedented, sustained transmission among mammals, indicate enhanced adaptation of the H5N1 clade 2.3.4.4b virus to mammalian hosts (14, 15). To date, 71 H5N1 human cases have been reported in the U.S., including one fatality, primarily linked to occupational exposure on dairy farms and poultry operations, or during culling activities in affected farms (16).

Natural infections with H5N1 genotype B3.13 viruses in dairy cattle have been associated with severe viral mastitis, characterized by lower milk quality and decreased milk production, with affected animals shedding high titers of infectious virus in milk (10, 14, 17). Experimental infections in lactating cows via intramammary inoculation successfully recapitulated these clinical signs (18, 19). A few critical mammalian-adaptive mutations, including E627K and M631L amino acid substitutions in the viral polymerase basic 2 (PB2) protein, have been associated with enhanced replication of H5N1 genotype B3.13 viruses in the mammary tissue (11, 19, 20). Notably, the virus characterized from one of the first H5N1 B3.13 human cases in Texas (Texas/37) harbored the E627K PB2 mutation (21). Previous studies have shown that the PB2 E627K, Q591R and D701N mutations help effective binding of the viral polymerase complex with mammalian acid nuclear phosphoprotein 32 (ANP32) proteins and thereby enable viruses with these mutations to use the mammalian ANP32 more effectively (22).

Despite these concerning developments, data on the pathogenicity and transmissibility of bovine- and human-derived H5N1 genotype B3.13 viruses in mammalian models remain limited (15, 23, 24). Previous studies have shown that H5N1 genotype B3.13 viruses cause lethal systemic infection in mice and ferrets, with limited evidence of transmission to naïve contact adult animals or to pups (15, 20, 24). Here, we investigated the pathogenesis and contact transmission potential of H5N1 genotype B3.13 viruses derived from bovine and human hosts in murine and Syrian hamster models of infection. We evaluated and compared viral replication kinetics, tissue tropism, and resulting pathology, clinical outcomes, and transmissibility. Our findings provide critical insights into the pathogenesis of the emergent H5N1 genotype B3.13 virus in relevant mammalian models of infection.

## Results

### Characterization of recombinant H5N1 clade 2.3.4.4b genotype B3.13 viruses derived from bovine and human hosts

Using reverse genetics, we rescued three naturally occurring H5N1 viruses of the clade 2.3.4.4b genotype B3.13, including a bovine-derived virus (rTX2/24) and two human-derived viruses (rTexas/37 and rMichigan/90). The reverse genetics system was first established using the A/Cattle/Texas/063224-24-1/2024 virus isolate (TX2/24; GISAID accession: EPI_ISL_19155861) as a backbone. Full-length gene segments were synthesized and cloned into the bidirectional pHW2000 vector. The recombinant TX2/24 virus (rTX2/24) was successfully rescued by transfecting a co-culture of human embryonic kidney 293T (HEK293T) and bovine uterine epithelial (Cal-1) cells.

To rescue the human-derived viruses, we compared the genome sequences of two human H5N1 B3.13 isolates - A/Texas/37/2024 (EPI_ISL_19027114) and A/Michigan/90/2024 (EPI_ISL_19162802) - with that of the bovine A/Cattle/Texas/063224-24-1/2024 isolate (Fig. 1A). Gene segments that differed between the bovine and human-derived virus were replaced, including PB2 (G362E, E627K and L631M), PB1 (I392V), PA (K142E, I219L, R497K), and NS (Q40R) from Texas/37-, and PB1 (E581K and A741V), NA (Q51R), and NS (L27M) from the Michigan/90 viruses. These segments were synthesized to match the respective genome segments of human Texas/37 and Michigan/90, cloned into the pHW2000 vector, and used to rescue the rTexas/37 and rMichigan/90 viruses via transfection of HEK293T-Cal-1 co-cultures. Working stocks of all recombinant viruses were prepared in 9- to 10-day-old embryonated chicken eggs. Whole-genome sequencing confirmed the identity of the recombinant viruses to their parental bovine (TX2/24) or human origin (Texas/37 and Michigan/90) viruses and verified the absence of unintended mutations.

**FIG 1.**
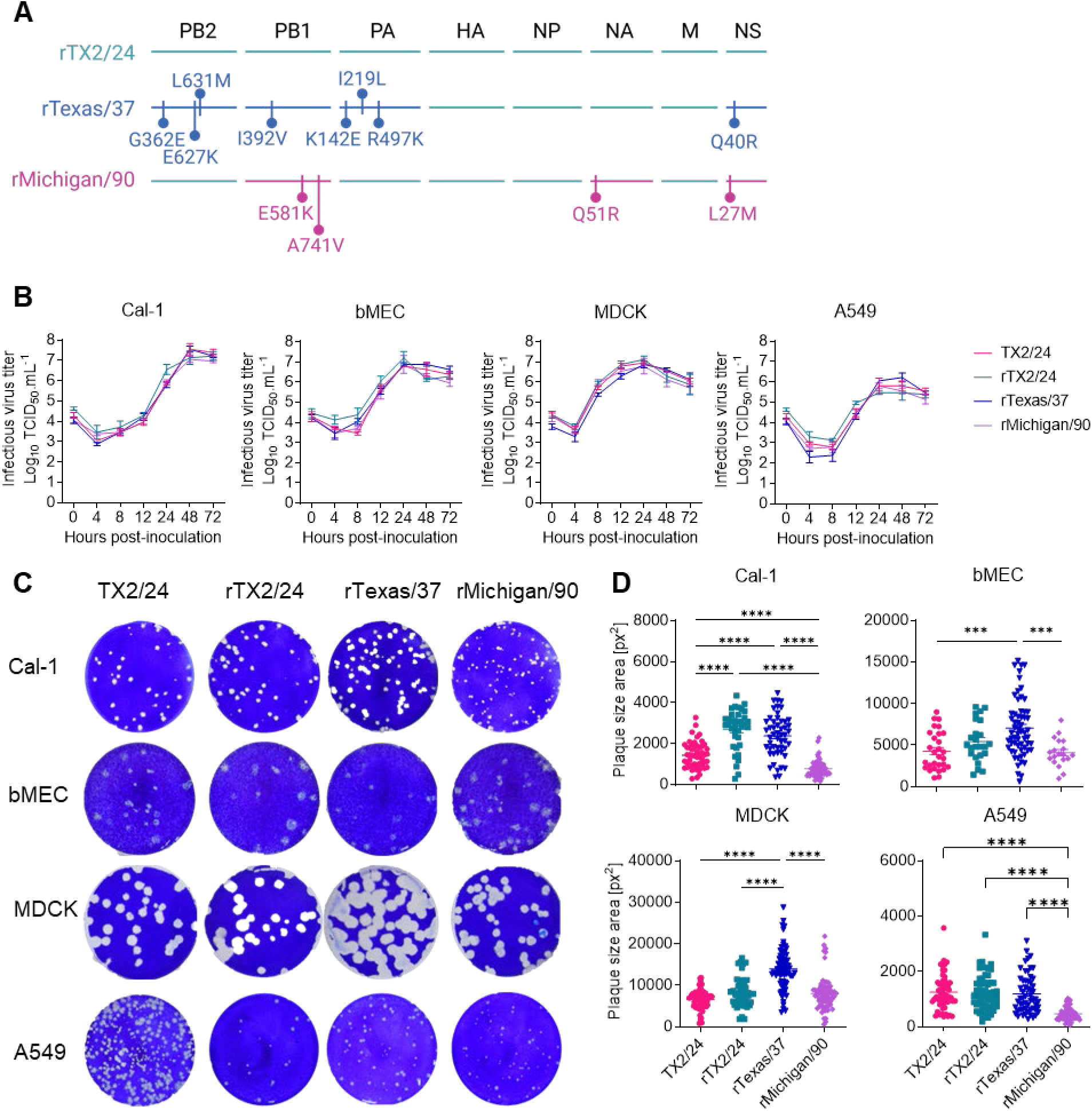
Replication kinetics and plaque size phenotype of HPAI H5N1 viruses in mammalian cells. (A) Genomic schematics of recombinant viruses based on bovine TX2/24 and human Texas/37 and Michigan/90 gene sequences. Amino acid differences between the bovine TX2/24 virus and the human-derived Texas/37 and Michigan/90 are shown. (B) Multicycle growth curves. Bovine uterine epithelial cells (Cal-1), primary bovine mammary epithelial cells (bMEC), Madin-Darby canine kidney (MDCK) and human lung adenocarcinoma (A549) cells were infected (MOI 0.1) with HPAI H5N1 TX2/24, rTX2/24, rTexas/37 and rMichigan/90 viruses and virus titers were determined at indicated time points by limiting dilution method and expressed as TCID50.mL^-1^. Data indicates mean ± SEM, n = 3, three independent experiments. (C) Viral plaque phenotypes. Cal-1, bMEC, MDCK and A549 cells cultured in 6-well plates were infected (30 plaque-forming unit/well) with HPAI H5N1 isolates and overlaid with medium containing 1% agar. Plates were incubated at 37 °C for 72 h, the agar overlay was removed, cells were fixed with 3.7% formaldehyde and the monolayer was stained with 0.5% crystal violet. (D) The diameters of viral plaques from the pictures in C were measured using ViralPlaque software in pixel^2^ and plotted. One-way ANOVA with Tukey’s multiple comparison test, *** *p* < 0.001 and **** *p* < 0.0001.

We next characterized the wild-type and recombinant viruses *in vitro* by assessing their antiviral sensitivity, replication kinetics and plaque size phenotypes (Fig. 1B-1D, Supplementary Figure 1). Antiviral sensitivity analysis using two common influenza A antiviral drugs, oseltamivir (neuraminidase inhibitor) and amantadine (M2 proton channel inhibitor), revealed comparable susceptibility of the three rescued recombinant viruses to these two antivirals *in vitro* (Supplementary Fig. 1). Multi-step growth curves were performed in four mammalian cell lines, including Cal-1, primary bovine mammary epithelial cells (bMEC), Madin-Darby canine kidney (MDCK), and human lung adenocarcinoma (A549) cells. The wild-type TX2/24 virus and three recombinant viruses of bovine and human origin (rTX2/24, rTexas/37, and rMichigan/90) exhibited comparable replication efficiency across all cell types (Fig. 1B). Plaque size analysis revealed both cell-type and virus-specific differences (Fig. 1C-1D). MDCK and bMEC cells consistently supported the formation of larger plaques compared to Cal-1 and A549 cells. The wild-type TX2/24 and its recombinant counterpart rTX2/24 produced similar plaque sizes in most cell lines, except for a notable difference with smaller plaques produced by the rTX2/24 virus in Cal-1 cells. Interestingly, the human-derived rTexas/37 virus formed significantly larger plaques (p ≤ 0.001) across most cell types compared to both rTX2/24 and rMichigan/90. In contrast, the rMichigan/90 virus consistently produced smaller plaques (p ≤ 0.001) in all cell lines compared with both rTX2/24 and rTexas/37 (Fig. 1C-1D). Collectively, wild-type and recombinant H5N1 viruses showed similar replication kinetics across cell lines, but exhibited distinct, virus- and cell-type-dependent plaque size phenotypes.

### Polymerase activity of bovine- and human-derived H5N1 clade 2.3.4.4b viruses

We compared the polymerase activity of the TX2/24, Texas/37, and Michigan/90 viruses using *in vitro* replicon assays. Individual viral polymerase subunits (PB2, PB1, and PA) and NP from each virus were cloned into mammalian expression plasmids (pCAGGS) and co-transfected into HEK293T cells with either the pUC57-NP(NCR)-NLuc or pUC57-NP(NCR)-miniGFP2 minigenome plasmid. A negative control, consisting of HEK293T cells co-transfected with PB2-, PA- and NP-expressing plasmids (lacking PB1) plus the minigenome plasmids were included. At 24 h post-transfection, the human-derived Texas/37 virus exhibited significantly higher polymerase activity than the bovine-derived TX2/24 and human-derived Michigan/90 viruses, as indicated by stronger GFP expression and increased luminescence (*p* ≤ 0.01) (Fig. 2A and B). In contrast, polymerase activities of the three viruses were comparable at 48 h post-transfection, with minimal differences in GFP expression and luminescence (Fig. 2C). No GFP expression or luminescence was detected in the absence of the PB1 polymerase subunit in negative control cells.

**FIG 2.**
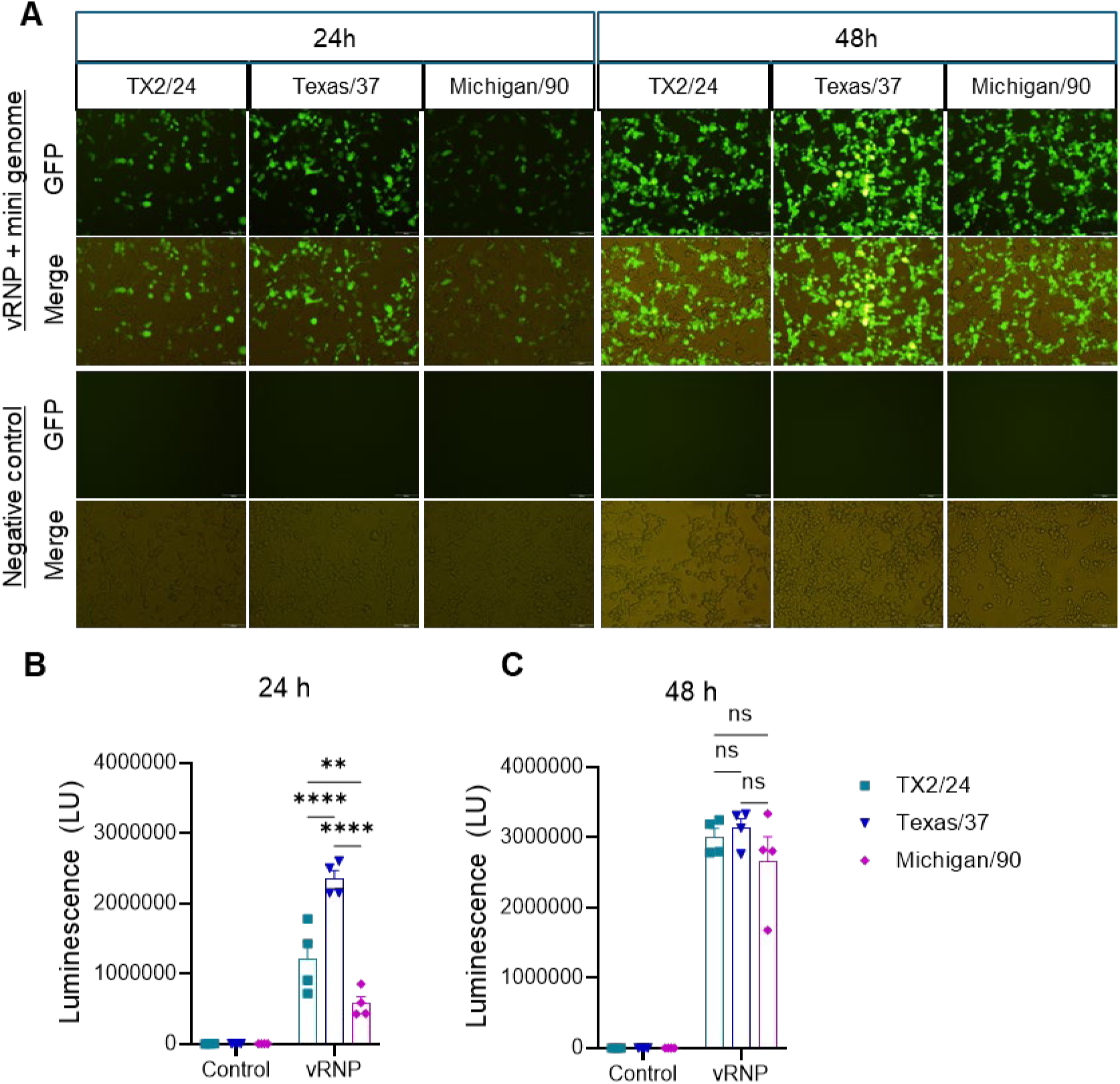
Polymerase activity of H5N1 clade 2.3.4.4b viruses. HEK293T cells were co-transfected with pcAGGS plasmids encoding the viral ribonucleoprotein (RNP) complex components PB2, PA, and NP, with or without PB1, with either the pUC57-NP(NCR)-miniGFP2 (A) or pUC57-NP(NCR)-NLuc (B–C) minigenome reporter plasmids. GFP expression (A) and nanoluciferase activity (B-C) were analyzed by fluorescence microscopy and luciferase reporter assay, respectively. Data represents the mean ± SEM from four independent experiments.

### Bovine- and human-derived H5N1 clade 2.3.4.4b genotype B3.13 viruses cause lethal infections in mice

To compare the pathogenicity of the H5N1 viruses studied here, we inoculated C57BL/6 mice intranasally with 1x10^3^ PFU of bovine-derived TX2/24, recombinant rTX2/24, and human-derived rTexas/37 or rMichigan/90 viruses. Mice were monitored daily for five days and euthanized once reaching pre-determined humane endpoints (≥20% body weight loss) (Fig. 3A). Mice infected with TX2/24 (5/6), rTX2/24 (6/6), and rTexas/37 (6/6) exhibited rapid weight loss and reached humane endpoints by day 4 post-infection (pi). The remaining mouse in the TX2/24 group and all mice infected with rMichigan/90 reached humane endpoints by day 5 pi (Fig. 3B). In contrast, mice in the mock-infected control group maintained steady body weight throughout the study. Daily mortality is shown in the survival curves (Fig. 3C).

**FIG 3.**
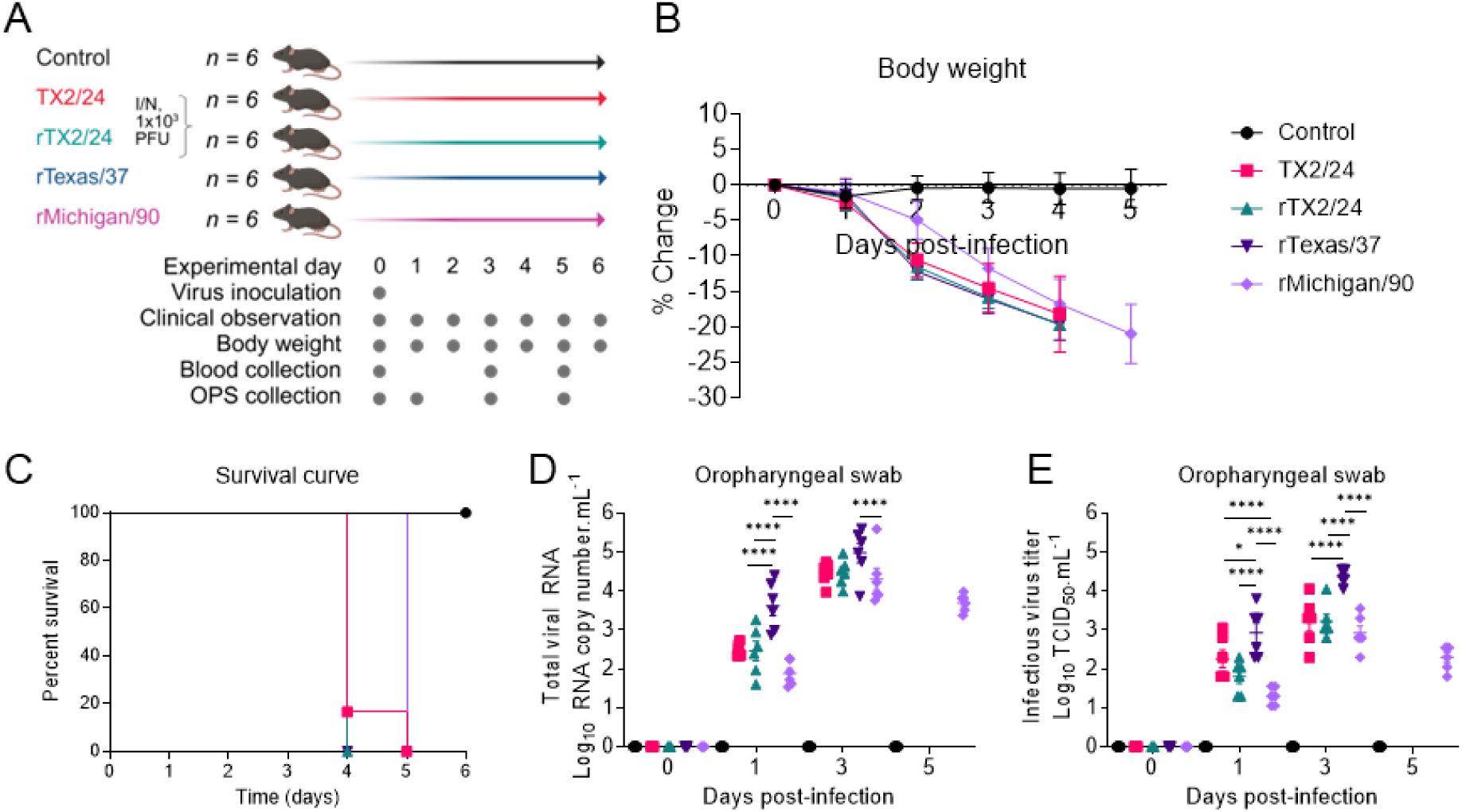
Pathogenesis of bovine- and human-derived HPAI H5N1 viruses in mice. (A) Experimental design. (B) C57BL/6 mice (*n* = 30) were mock inoculated (*n* = 6) or inoculated intranasally with 1x10^3^ PFU of HPAI H5N1 TX2/24 (*n* = 6), rTX2/24 (*n* = 6), rTexas/37 (*n* = 6) and rMichigan/90 (*n* = 6) viruses. Animals were monitored and body weight was recorded daily for 5 days post-infection (pi). (C) Survival curves. H5N1 viral RNA (D) and infectious viral (E) loads quantified by rRT-PCR and virus titration in oral secretions collected on days 0, 1, 3 and 5 pi. Virus titers were determined by endpoint dilution and expressed as TCID_50_.mL^-1^. The limit of detection (LOD) for infectious virus titration was 10^1.05^ TCID_50_.mL^-1^. Two-way ANOVA with Tukey’s multiple comparison test, * *p* < 0.05 and **** *p* < 0.0001.

We next quantified viral RNA and infectious virus titers in oropharyngeal swabs collected on days 0, 1, 3, and 5 dpi using rRT-PCR and virus titration in Cal-1 cells, respectively (Figs. 3D-E). Mice infected with the TX2/24 and rTX2/24 viruses showed comparable levels of viral RNA and infectious titers throughout the study (Fig. 3D-E). Notably, mice infected with rTexas/37 showed significantly higher (*p* ≤ 0.05) viral RNA (3.62 ± 0.62 log_10_ genome copies.mL^-1^) loads on day 1 compared to mice infected with TX2/24 (2.48 ± 0.17 log_10_ genome copies.mL^-1^), rTX2/24 (2.46 ± 0.61 log_10_ genome copies.mL^-1^), and rMichigan/90 (1.88 ± 0.32 log_10_ genome copies.mL^-1^). Viral RNA levels in oral secretions increased on day 3 pi and were comparable between mice inoculated with TX2/24, rTX/24 and rTexas/37. However, rMichigan/90-inoculated mice showed significantly (*p* ≤ 0.001) lower viral RNA load than animals inoculated with rTexas/37 (Fig. 3D). Similar to the viral RNA loads, infectious virus titers were also significantly higher (*p* ≤ 0.05) in mice inoculated with rTexas/37 (2.93 ± 0.63 log_10_ TCID_50_.mL^-1^) on day 1 pi when compared to the bovine TX2/24 (2.26 ± 0.56 log_10_ TCID_50_.mL^-1^) and rTX2/24 (1.8 ± 0.42 log_10_ TCID_50_.mL^-1^) and human rMichigan/90 (1.3 ± 0.22 log_10_ TCID_50_.mL^-1^) inoculated groups (Fig. 3E). Among all groups, mice infected with rMichigan/90 exhibited the lowest infectious virus titers on day 1 pi. On day 3 pi, the infectious virus titers increased in all inoculated groups, with the rTexas/37 inoculated mice showing the highest virus loads (4.38 ± 0.2 log_10_ TCID_50_.mL^-1^) in the oral secretion (Fig. 3E). No viral RNA or infectious virus was detected in mock-inoculated mice throughout the experiment. Taken together, both bovine- and human-derived H5N1 clade 2.3.4.4b genotype B3.13 viruses caused lethal disease in mice, with rMichigan/90 showing slightly decreased virulence and reduced viral replication and shedding compared to the other H5N1 genotype B3.13 viruses in our study.

### Virus replication and tissue tropism in mice infected with HPAI H5N1 viruses

We quantified HPAI viral RNA (Fig. 4A) and determined infectious viral (Fig. 4B) loads in brain, trachea, lungs, liver, spleen, small intestine, and large intestine of mice by rRT-PCR and virus titrations, respectively. Viral RNA loads were highest in the brain of TX2/24-infected mice (6.75 ± 0.27 log_10_ genome copies.g^-1^) compared with rTX2/24- (4.62 ± 0.46 log_10_ genome copies.g^-1^), rTexas/37- (5.49 ± 0.33 log_10_ genome copies.g^-1^) and rMichigan/90- (4.86 ± 0.61 log_10_ genome copies.g^-1^) groups. Similar trends were observed in trachea and liver, with relatively higher RNA loads in TX2/24-infected mice. In contrast, lungs exhibited the highest viral RNA loads overall (7.25-7.87 log_10_ genome copies.g^-1^), which were comparable among the four infected groups. Notably, rMichigan/90-infected mice had lower RNA loads in trachea, liver, and spleen compared with TX2/24 (Fig. 4A). Infectious virus titration revealed higher titers in the brain (4.17 ± 0.43 log_10_ TCID_50_.g^-1^) and liver (5.38 ± 0.49 log_10_ TCID_50_.g^-1^) of TX2/24-infected mice relative to the other groups (Fig. 4B). The lungs had the highest infectious virus titers across tissues (5.67-6.39 log_10_ TCID_50_.g^-1^), with no significant differences between groups. Virus titers in trachea and spleen were also comparable among infected mice. Viral RNA, but no infectious virus, was detected in small and large intestines. No HPAI RNA or infectious virus was detected in mock-inoculated mice.

**FIG 4.**
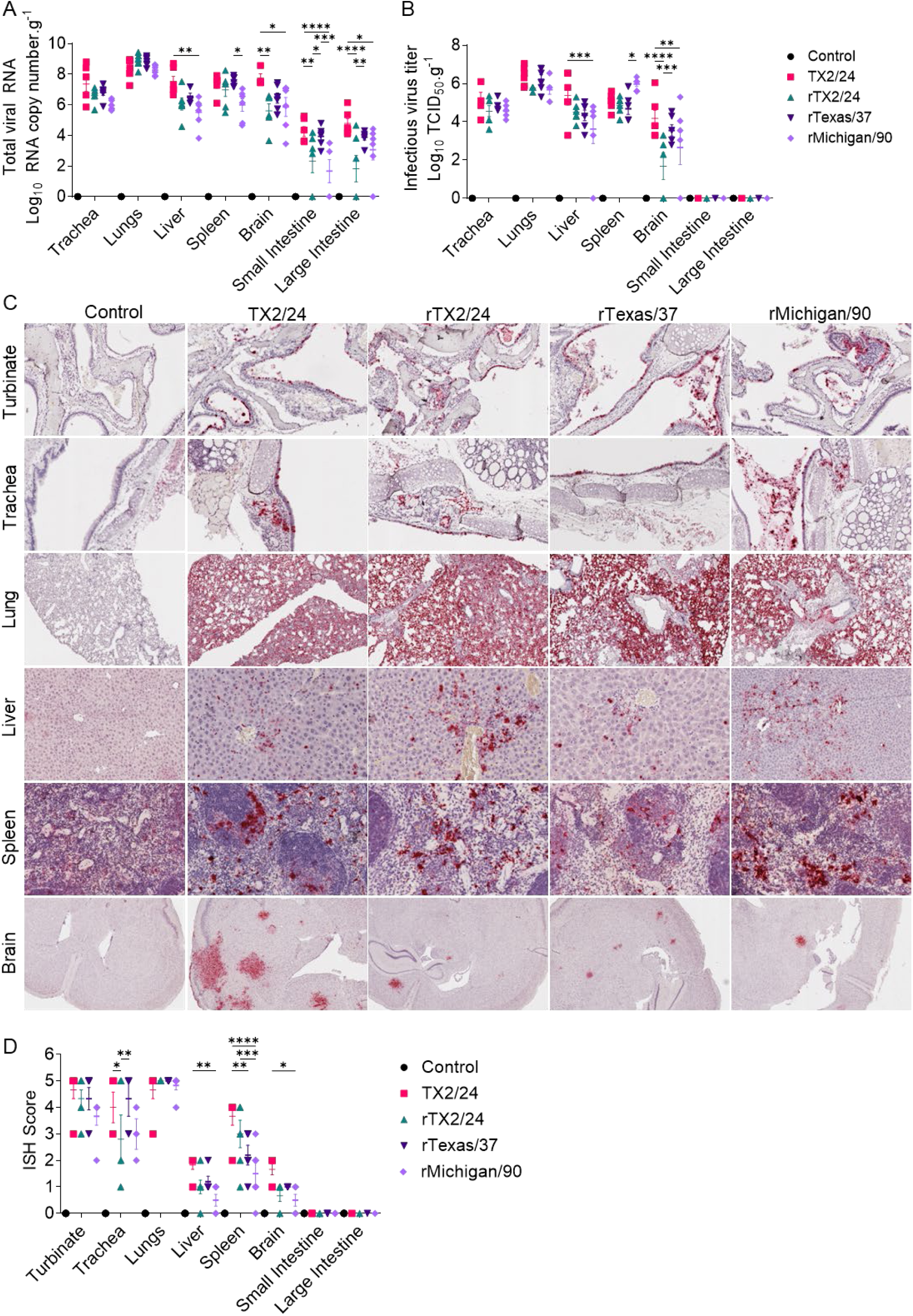
Virus distribution in tissues of mice infected with bovine- and human-derived HPAI H5N1 viruses. HPAI H5N1 viral RNA (A) and infectious virus (B) loads quantified by rRT-PCR and virus titration in tissues of mice collected at necropsy, respectively. *In situ* hybridization (ISH) scoring signal (C, red color) of viral RNA in tissues of HPAI H5N1 infected mice. ISH scoring of tissues from (C). Virus titers were determined using endpoint dilutions and expressed as TCID_50_.mL^-1^. The limit of detection (LOD) for infectious virus titration was 10^1.05^ TCID_50_.mL^-1^. (A-C) Data indicate mean ± SEM of 6 mice per group per tissues. Two-way ANOVA with Tukey’s multiple comparison test, * *p* < 0.05, ** *p* < 0.01, *** *p* < 0.001, and **** *p* < 0.0001.

Viral RNA localization was assessed by *in situ* hybridization (ISH) and histological lesions were scored (Fig. 4C-D, Supplementary Table 1) (25). In the brain, all viruses caused diffuse and multifocal hybridization signal, with TX2/24 producing more foci and higher lesion scores. The hybridization signal was concentrated on neurons around blood vessels in the peri-ventricular areas. In nasal turbinates, intense hybridization signals were observed in the epithelial lining and exudates, with comparable RNA levels among all groups. In the trachea, viral RNA was detected in the epithelial lining, intraluminal sloughed material, and submucosal glands, with slightly lower scores in rMichigan/90-infected mice. In the lungs, an extensive hybridization signal was present in alveolar epithelial cells, interalveolar septa, and bronchial/bronchiolar epithelium, with similar intensities across all groups. In the liver, a sparse hybridization signal occurred in hepatocytes and sinusoids near portal areas, with slightly lower scores in rMichigan/90. In the spleen, viral RNA hybridization signal was mainly observed in the red pulp with sparse hybridization signal in germinal centers, again with slightly lower scores in rMichigan/90-infected animals. Both small and large intestine epithelium exhibited minimal hybridization signal. Taken together, the TX2/24- infected mice displayed higher viral loads and broader tissue distribution, particularly higher viral loads in the brain and liver, whereas the rMichigan/90-infected mice generally had lower viral RNA levels and lesion scores.

Histopathologic changes were mostly restricted to the lung and liver, with minimal lesions noted in the nasal turbinates and trachea (Supplementary Figure 2, Supplementary Table 1). Mild to moderate lymphocytic and neutrophilic interstitial pneumonia was noted in the lung with occasional alveolar necrosis and fibrin accumulation. Changes in the liver were characterized by mild to moderate lymphocytic and neutrophilic hepatitis with hepatocellular necrosis. Mild lymphocytic inflammation was noted in the trachea and nasal turbinates with rare areas of erosion and ulceration.

### Bovine- and human-derived H5N1 clade 2.3.4.4b genotype B3.13 viruses caused a milder acute infection in hamsters

We evaluated the pathogenicity of bovine- and human-derived H5N1 clade 2.3.4.4b genotype B3.13 viruses in Golden Syrian hamsters (n = 39). Animals were either mock inoculated (*n* = 3) or intranasally inoculated with 5x10^5^ PFU of TX2/24 (bovine, *n* = 9), rTX2/24 (recombinant TX2/24, *n* = 9), rTexas/37 (human, *n* = 9), or rMichigan/90 (human, *n* = 9). Three animals from each group were euthanized on days 3, 5, and 7 post-infection (pi); mock-inoculated animals were euthanized on day 7 (Fig. 5A). Hamsters infected with TX2/24 and rTX2/24 exhibited gradual weight loss, reaching 7.86% and 10.45% reductions by day 7 pi, respectively. Similarly, rMichigan/90-infected hamsters showed progressive weight loss, reaching an 8.73% reduction in body weight by day 7. In contrast, rTexas/37-infected hamsters experienced a rapid weight loss (7.76%) by day 2 pi, which was maintained until day 5, when the animals recovered and regained body weight until day 7 (Fig. 5B).

**FIG 5.**
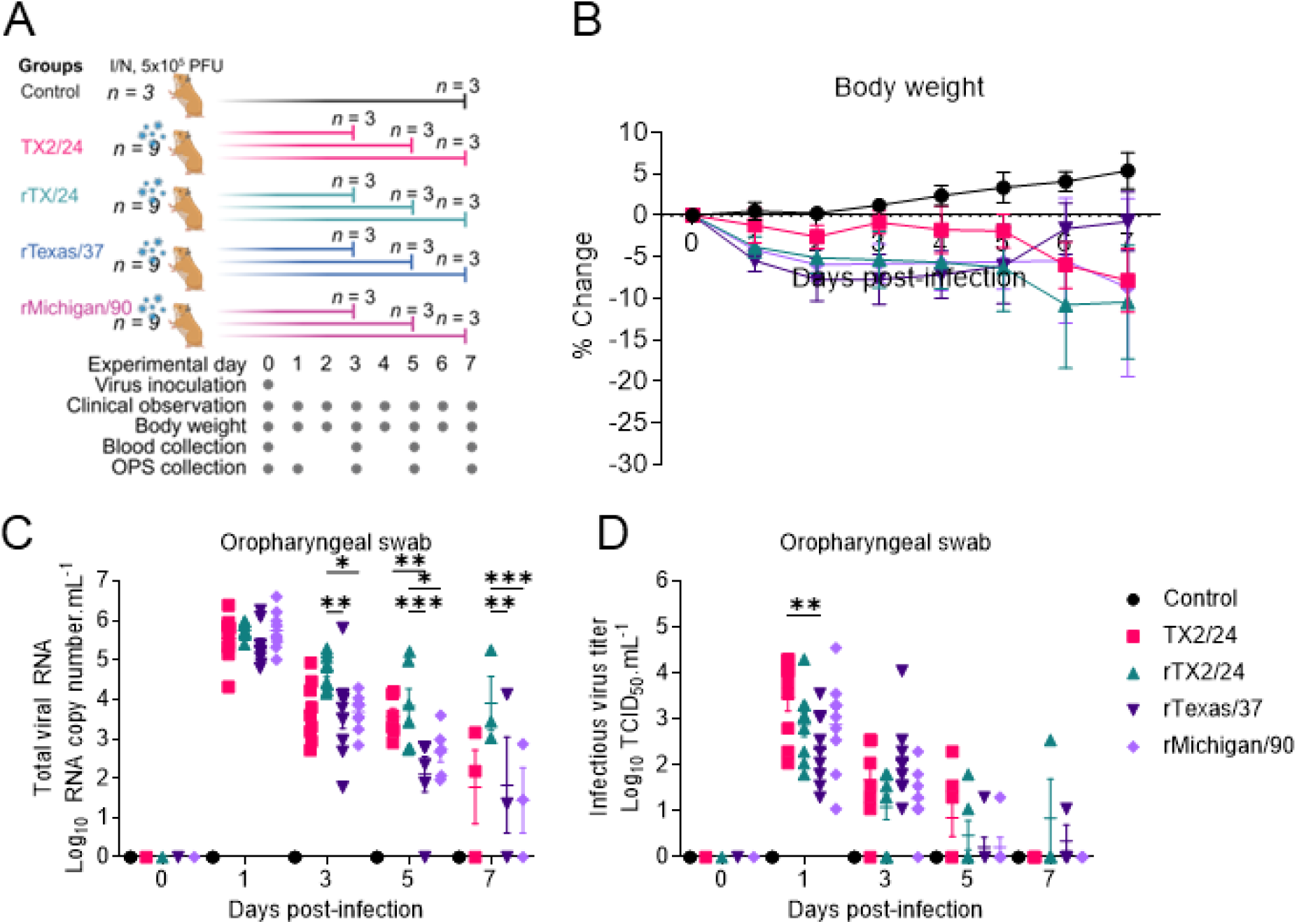
Pathogenesis of bovine- and human-derived HPAI H5N1 viruses in hamsters. (A) Experimental design. (B) Golden Syrian hamsters (*n* = 39) were mock inoculated (*n* = 3) or inoculated intranasally with 5x10^5^ PFU of HPAI H5N1 TX2/24 (*n* = 9), rTX2/24 (*n* = 9), rTexas/37 (*n* = 9) and rMichigan/90 (*n* = 9) viruses and changes in body weight gain were monitored for 7 days post-infection (pi). HPAI viral RNA (C) and infectious viral (D) loads quantified by rRT-PCR and virus titration, respectively in oral secretions collected on days 0, 1, 3, 5 and 7 pi. Virus titers were determined using endpoint dilutions and expressed as TCID_50_.mL^-1^. The limit of detection (LOD) for infectious virus titration was 10^1.05^ TCID_50_.mL^-1^. Two-way ANOVA with Tukey’s multiple comparison test, * *p* < 0.05, ** *p* < 0.01 and *** *p* < 0.001.

To assess viral shedding, we quantified H5N1 viral RNA and infectious virus titers in oral secretions collected from infected hamsters on days 0, 1, 3, 5, and 7 pi. The highest viral RNA levels were observed on day 1 pi, with comparable viral loads observed across all infected groups (5.37-5.74 log_10_ genome copies.mL^-1^) (Fig. 5C). By day 3 pi, viral RNA levels declined by 1-2 log_10_, with significantly lower levels detected in hamsters infected with human-derived rTexas/37 (3.64 ± 1.13 log_10_ genome copies.mL^-1^) and rMichigan/90 (3.68 ± 0.44 log_10_ genome copies.mL^-1^) compared to bovine-derived rTX2/24 (4.72 ± 0.43 log_10_ genome copies.mL^-1^) (*p* ≤ 0.05). This trend persisted on days 5 and 7 pi, with both human virus-infected groups shedding significantly less viral RNA than the bovine virus group.

Infectious virus titers in oral secretions followed a similar pattern. On day 1 pi, rTexas/37- infected hamsters exhibited significantly lower titers than those infected with TX2/24 (*p* ≤ 0.01) (Fig. 5D), while titers in rMichigan/90-, TX2/24-, and rTX2/24-infected groups were comparable. By days 3, 5, and 7 pi, infectious virus levels decreased gradually and did not significantly differ among the four groups. Collectively, the bovine-derived H5N1 B3.13 viruses showed higher and more sustained replication and viral shedding in hamsters compared to human-derived strains.

### Virus replication and tissue tropism in hamsters infected with HPAI H5N1 viruses

We quantified the distribution of viral RNA and infectious viruses in tissues of hamsters infected with bovine- and human-derived H5N1 isolates (Supplementary Fig. 3). Tissue samples (mandibular lymph node, olfactory bulb, brain, nasal turbinate, trachea, lung, heart, stomach, pancreas, intestine, liver, spleen, kidney, testes, ovary, and uterus) were collected at necropsy on days 3, 5, and 7 post-infection (pi) and analyzed by rRT-PCR and virus titration.

On day 3 pi, the highest viral RNA loads were detected in the nasal turbinate (6.64-7.71 log_10_ genome copies.g^-1^) and lung (4.26-6.65 log_10_ genome copies.g^-1^), with comparable levels across the four inoculated groups (Supplementary Fig. 3A). Intermediate viral RNA levels were observed in the olfactory bulb, which were significantly higher in the bovine-derived TX2/24 group (4.57-6.25 log_10_ genome copies.g^-1^) compared to the human-derived rTexas/37 (3.53 ± 2.26 log_10_ genome copies.g^-1^) and rMichigan/90 (2.09 ± 1.05 log_10_ genome copies.g^-1^) groups. Low levels of viral RNA were detected in the brain, heart, stomach, intestine, liver, spleen, kidney, and reproductive organs (Supplementary Fig. 3A).

By day 5 pi, viral RNA loads in the lungs declined markedly (1.6-4.06 log_10_ genome copies.g^-1^) in all groups, although rTexas/37-infected hamsters showed relatively higher levels than the others (Supplementary Fig. 3C). Viral RNA in the nasal turbinate remained comparable across groups (5.5-6.81 log_10_ genome copies.g^-1^), showing an approximately 1-log reduction from day 3. In contrast, viral RNA loads in the olfactory bulb remained significantly higher (p ≤ 0.001) in TX2/24 (5.19 ± 1.44 log_10_ genome copies.g^-1^) and rTX2/24 (5.97 ± 1.33 log_10_ genome copies.g^-1^) than in the rTexas/37 (1.2 ± 0.6 log_10_ genome copies.g^-1^) and rMichigan/90 (1.71 ± 0.08 log_10_ genome copies.g^-1^) groups (Supplementary Fig. 3C).

On day 7 pi, significantly higher viral RNA loads (p ≤ 0.05) were detected in the olfactory bulb and brain of TX2/24 (8.25 ± 0.16 and 8.23 ± 0.19 log_10_ genome copies.g^-1^) and rTX2/24 (7.95 ± 1.18 and 8.19 ± 0.89 log_10_ genome copies.g^-1^) groups, compared to rTexas/37 (undetected and 0.55 ± 0.55 log_10_ genome copies.g^-1^) and rMichigan/90 (4.46 ± 1.82 and 3.43 ± 2.44 log_10_ genome copies.g^-1^) (Supplementary Fig. 3E). Sporadic viral RNA was detected in the trachea and lungs on this day.

Virus titration confirmed infectious viruses in the olfactory bulb, brain, nasal turbinate, trachea, and lung (Supplementary Fig. 3B, D, E). In nasal turbinates, comparable infectious virus titers were detected on day 3 (3.13-3.97 log_10_ TCID_50_.g^-1^) and day 5 (2.7-3.38 log_10_ TCID_50_.g^-1^) across all groups, but no infectious virus was detected in rTexas/37-inoculated animals on day 7. Infectious viruses were mainly detected in the lungs on day 3, with comparable titers across groups. In contrast, the viruses were predominantly cleared from the olfactory bulb and brain on day 7 in TX2/24- and rTX2/24-infected animals, while little to no infectious virus was detected in rTexas/37- and rMichigan/90-infected hamsters.

Viral RNA distribution was assessed by *in situ* hybridization (ISH) in tissues collected at necropsy on day 3, 5 and 7 pi (Fig. 6). Among all tissues tested, viral RNA hybridization signal was observed only in the lungs and brain. Extensive hybridization signal was present in alveolar epithelial cells, interalveolar septa, and bronchial/bronchiolar epithelium on day 3 pi, with slightly higher intensities and number of infected cells in rTX2/24, rTexas/37 and rMichigan/90-inoculated animals than the TX2/24-inoculated ones (Fig. 6A). No RNA hybridization signal was detected in the brain of hamsters inoculated with both bovine- and human-derived viruses on days 3 and 5 pi, except the TX2/24 in which one hamster showed hybridization signal (Fig. 6B). Intense viral RNA hybridization signal spreading to considerable areas of the brain was observed on day 7 in hamsters inoculated with TX2/24, rTX2/24 and rMichigan/90 viruses. No viral RNA hybridization signal in the brain was detected in any of the hamsters inoculated with rTexas/37 on day 7. Viral RNA hybridization signal was mostly detected in neurons as well as in glial cells and ependymal cells lining the ventricles. Collectively, the bovine-derived H5N1 isolates (TX2/24 and rTX2/24) replicated more efficiently in the olfactory bulb and brain of hamsters than human-derived isolates (rTexas/37 and rMichigan/90), which remained largely restricted to the respiratory tract during a short time frame.

**FIG 6.**
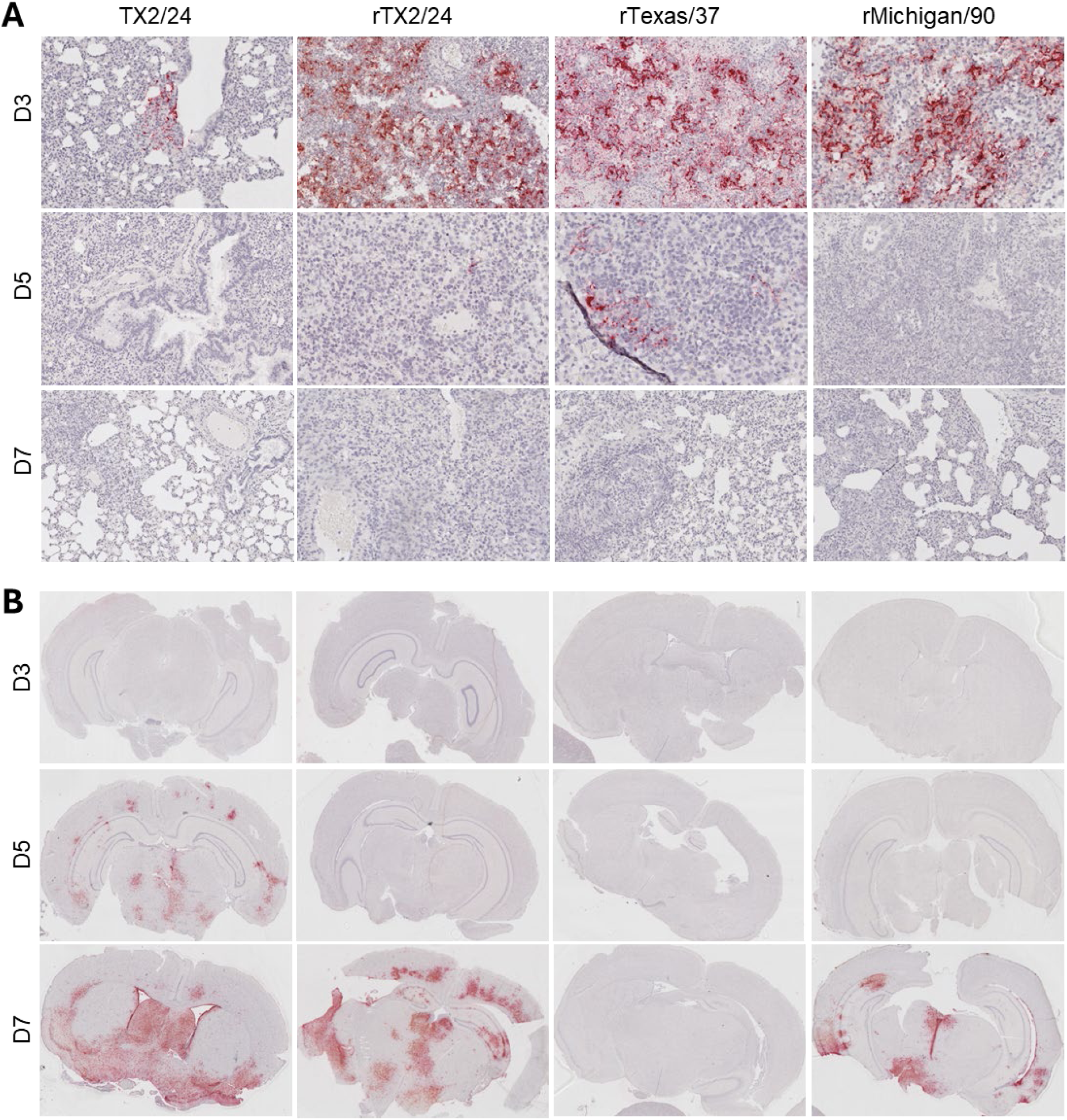
*In situ* distribution of HPAI H5N1 viral RNA in lung (A) and brain (B) of hamsters inoculated with bovine- and human-derived clade 2.3.4.4b viruses. Lung and brain tissues of hamsters were collected at necropsy on days 3, 5 and 7 pi and processed by *in situ* hybridization (ISH). Representative images of 3 animals per group per time point were shown.

Histopathologic changes were noted in the lungs and brain (Supplementary Figure 2 and Supplementary Table 2). In the lung, mild-to-severe lymphohistiocytic interstitial pneumonia was noted with evidence of alveolar necrosis and fibrin. Mild to severe encephalitis, lymphohistiocytic with occasional neutrophils, was noted along with neuronal necrosis, neuronophagia, satellitosis, and gliosis.

### Transmission potential of bovine- and human-derived H5N1 clade 2.3.4.4b genotype B3.13 viruses in hamsters

We evaluated the transmission potential of bovine- and human-derived clade 2.3.4.4b viruses in hamsters. Hamsters (n = 4/virus) were intranasally inoculated with 5 x 10^5^ PFU of TX2/24, rTX2/24, rTexas/37, or rMichigan/90 viruses. At 24 h post-inoculation (pi), each infected animal was co-housed with a naïve contact hamster (n = 4/virus) for 48 h (Fig. 5A). Body weight was recorded daily, oropharyngeal swabs (OPS) were collected daily between days 0-8, and then on day 10, and 14 pi to assess virus replication and shedding. Blood samples were obtained on days 0, 7, and 14 to assess seroconversion in both infected and contact animals (Fig. 7A).

**FIG 7.**
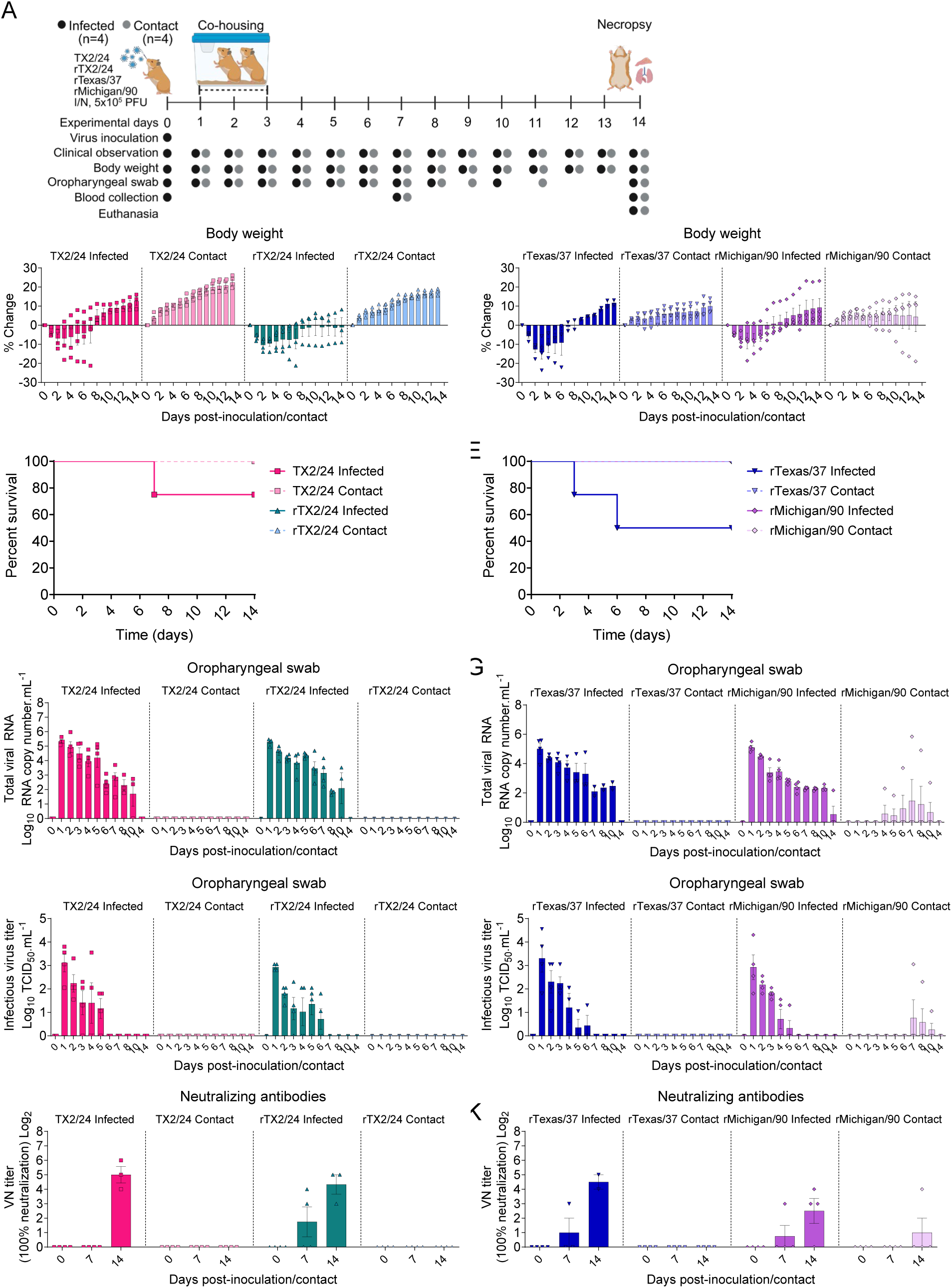
Contact transmission of bovine- and human-derived HPAI H5N1 viruses in hamsters. (A) Experimental design. (B-C) Changes in body weight of hamsters following intranasal inoculation of HPAI H5N1 TX2/24, rTX2/24, rTexas/37 and rMichigan/90 viruses and in contact animals throughout the 14-day experimental period. (D-E) Survival curves. (F-G) HPAI viral RNA load in oropharyngeal swabs quantified by rRT-PCR. (H-I) Infectious virus loads in oropharyngeal swabs determined by using endpoint dilution and expressed as TCID_50_.mL^-1^. The limit of detection (LOD) for infectious virus titration was 10^1.05^ TCID_50_.mL^-1^. (J-K) Neutralizing antibody responses to HPAI assessed by virus neutralization assay (100% neutralization) in serum. Two-way ANOVA with multiple comparison test, * *p* < 0.05 and ** *p* < 0.01.

Hamsters inoculated with bovine-derived TX2/24 and rTX2/24 began losing weight by day 1 pi, with loss continuing until days 2-4 pi, followed by recovery. However, one animal from each group continued to lose weight, reaching the humane endpoint (≥20% loss) on day 7 pi (Fig. 7B, 7D). The contact animals from these groups gained weight throughout the 14-day study period. Human-derived rTexas/37 and rMichigan/90 viruses produced similar early weight loss in inoculated animals. Two rTexas/37-infected hamsters reached humane endpoints on days 3 and 6 pi (Fig. 7C, 7E). All rMichigan/90-infected hamsters recovered after 6-8 days pi and gained weight thereafter. All contact animals from the rTexas/37 group gained weight throughout the study. In the rMichigan/90 group, all contact animals gained weight until day 4 post-contact (pc), after which one animal began losing weight, reaching ∼19% loss by the end of the study (Fig. 7C).

We analyzed the dynamics of HPAI viral replication and infectious virus shedding in the oropharyngeal swab (OPS) samples collected from both infected and contact hamsters (Fig. 7F- 7I). Viral RNA was consistently detectable in inoculated hamsters through day 8 pi, with sporadic detection on days 10 and 14 pi. Peak RNA loads occurred on day 1 pi (5.01-5.33 log_10_ genome copies.mL^-1^), followed by a gradual decline in RNA levels (Fig. 7F-7G). All contact animals from TX2/24, rTX2/24, and rTexas/37 groups tested negative by rRT-PCR. In contrast, one rMichigan/90 contact animal was RNA-positive starting on day 4 pc (2.29 log_10_ genome copies.mL^-1^), with peak viral RNA being detected at day 7 pc (5.84 log_10_ genome copies.mL^-1^). Infectious virus was detected in OPS from inoculated hamsters until days 5-6 pi, with peak viral titers on day 1 pi (2.92-3.3 log_10_ TCID_50_.mL^-1^) (Fig. 7H-7I). All contact animals from TX2/24, rTX2/24, and rTexas/37 groups were negative by virus isolation. The rMichigan/90 contact animal that tested positive for viral RNA also shed infectious virus from day 7 pc (3.05 log_10_ TCID_50_.mL^-1^) until day 10 pc (1.05 log_10_ TCID_50_.mL^-1^).

Neutralizing antibody titers measured by virus neutralization assay (Fig. 7J-7K) showed that by day 7 pi, two rTX2/24-infected hamsters were seropositive; by day 14 pi, all surviving TX2/24 (n = 3) and rTX2/24 (n = 3) animals had seroconverted. No contact animals in these groups seroconverted. In the rTexas/37 and rMichigan/90 groups, one infected hamsters from each group was seroconverted by day 7 pi, and all survivors seroconverted by day 14 pi. No contact animals in the rTexas/37 group seroconverted, whereas the contact animal in the rMichigan/90 group that shed virus seroconverted, confirming successful transmission of the rMichigan/90 virus to this animal. Overall, bovine- and human-derived clade 2.3.4.4b genotype B3.13 viruses showed limited transmission in hamsters, with only the rMichigan/90 virus transmitting to a single contact animal.

## Discussion

In this study, we characterized H5N1 clade 2.3.4.4b genotype B3.13 viruses derived from bovine (rTX2/24) and human (rTexas/37 and rMichigan/90) hosts, aiming at improving our understanding of their replication dynamics, pathogenicity, tissue tropism, and transmission potential in mice and/or hamster models of H5N1 virus infection. Our comparative studies in these relevant small animal models revealed that bovine-derived viruses disseminate more broadly, and present higher neurotropism in mice and hamsters, whereas human-derived viruses showed variable replication dynamics, ranging from enhanced replication and shedding (rTexas/37) to attenuated replication, yet transmitting to contact hamsters (rMichigan/90).

The differences observed between bovine- and human-derived viruses underscore potential host-specific adaptation. In mice, all viruses caused lethal disease, but the bovine TX2/24 exhibited the broadest tissue tropism, particularly targeting the brain, consistent with field reports of neuroinvasive properties and systemic disease in cattle, cats, and wild terrestrial mammals as well as with findings from experimental infections studies in mice and ferrets (2, 26–28). The mechanism of neuroinvasion of the bovine isolate remains undetermined, but establishment of viremia could favor hematogenous spread across the blood-brain barrier (26, 29), although dissemination of the virus through the olfactory route cannot be excluded. Notably, bovine-derived viruses also produced severe respiratory disease which were comparable to the two human-derived viruses. In contrast, the human-derived rTexas/37 virus replicated more efficiently in mice *in vivo* and produced higher early viral loads and larger plaques *in vitro*, suggesting enhanced polymerase activity favoring virus replication (30–32). Although the human-derived Texas/37 virus lacks the signature PB2 M631L substitution detected in all bovine-derived B3.13 H5N1 viruses, it encodes the mammalian-adaptive PB2 E627K mutation, which is known to facilitate interaction of the viral polymerase with mammalian ANP32 proteins, thereby enabling efficient replication of avian influenza viruses in mammals (30, 31). Consistent with this, replicon assays using miniGFP2 and nanoluciferase reporters confirmed higher polymerase activity for Texas/37 at early time points (24 h post-transfection) compared with TX2/24 and Michigan/90. At later time points (48 h), however, polymerase activity was comparable among all three viruses, suggesting that additional mammalian-adaptive mutations, including PB2 M631L, may contribute to enhanced polymerase function over time. Supporting this model, recent functional mapping of human- and bovine-derived H5N1 isolates further implicates PB2 residue 362 in the cap-binding domain: a G362E substitution modestly enhances polymerase activity and, importantly, can act synergistically with E627K to increase replication and pathogenicity in mice, supporting a model in which multiple PB2 changes jointly tune mammalian replication efficiency (32). Finally, the rMichigan/90 virus displayed a slightly attenuated phenotype, with smaller plaques across cell types *in vitro* and reduced virus replication and shedding in mice. This virus lacks the PB2 E627K mutation found in the Texas/37 virus but carries the bovine signature PB2 M631L substitution which is known to enhance the viral polymerase complex activity in mammalian cells (33, 34). The comparatively smaller plaque phenotype observed here suggests that additional substitutions, such as PB1 (E581K, A741V), NA (Q51R) or NS (L27M), may offset or modulate any replication advantage conferred by PB2 M631L.

Comparative analysis in hamsters revealed further divergence in tissue tropism. Bovine-derived viruses efficiently infected the olfactory bulb and brain, while human-derived isolates remained largely restricted to the respiratory tract. These results suggest that adaptation in bovines may enhance neuroinvasion, raising concerns for animal health, whereas human-derived strains may be undergoing early stages of adaptation toward sustained replication in the respiratory tract. Importantly, only the rMichigan/90 virus was transmitted, albeit inefficiently, as only a single contact hamster got infected. This apparent trade-off between virulence and transmissibility mirrors patterns described for influenza viruses adapting to new mammalian hosts. Our transmission study in hamsters corroborates a recent study in ferrets showing limited transmissibility of the bovine-derived H5N1 virus in ferrets (15). On the contrary, the US CDC study reported 100% contact transmission of the human-derived Texas/37 isolate in ferrets, which could be due to the differences in the host susceptibility of the virus between these two species (35). Similarly, Iwatsuki-Horimoto and colleagues also reported direct contact transmission of the human Texas/37 virus in a hamster model (36). These differences could be attributed to the differences in the experimental design, as Iwatsuki-Horimoto and colleagues co-housed the infected and contact hamsters throughout the study period (36), leading to increased opportunities for transmission, while we used a more stringent transmission model, maintaining the animals in contact for only 48 hours (35).

These findings carry important implications for zoonotic and pandemic risk. The ongoing detection of clade 2.3.4.4b viruses in avian and mammalian hosts and the continuous circulation of the virus in cattle with spillover to humans provide unique opportunities for virus adaptation. Our results show that while bovine-derived viruses are highly neurotropic and pathogenic, human-derived isolates may acquire phenotypic traits that favor transmissibility, even at the cost of reduced virulence. Together, these data underscore the urgent need for intensified surveillance of H5N1 viruses at the wildlife-livestock-human interface. Also critically important is the continuous functional characterization of emerging genotypes/strains/variants to improve our understanding of genotypes and phenotypes associated with mammalian adaptation.

## Materials and Methods

### Biosafety and biosecurity, and ethical regulations

All work involving rescuing, handling and propagation of HPAI H5N1 viruses was performed following strict biosafety measures in the Animal Health Diagnostic Center (AHDC) research BSL-3 suite at the College of Veterinary Medicine, Cornell University. The animal study procedures were reviewed and approved by the Institutional Animal Care and Use Committee at Cornell University (IACUC approval number 2024-0094). All relevant ethical regulations were followed.

### Cells

Bovine uterine epithelial cells (Cal-1, developed in house at the Virology Laboratory at AHDC) and Madin-Darby canine kidney (MDCK) cells were cultured in minimal essential medium (MEM, Corning Inc., Corning, NY) supplemented with 10% fetal bovine serum (FBS) and penicillin-streptomycin (ThermoFisher Scientific, Waltham, MA; 10 U.mL^–1^ and 100 µg.mL^–1^, respectively). Primary bovine mammary epithelial cells (bMEC; kindly provided by Dr. Sabine Mann, Department of Population Medicine and Diagnostic Sciences, College of Veterinary Medicine, Cornell University) were grown in William’s Medium E supplemented with 10% FBS, L-glutamine (Gibco, U.S.), 1X Insulin-Transferrin-Selenium (ITS) (Gibco, U.S.) and 10 ng/mL Epithelial Growth Factor (EGF) (VWR - Life Cell Technology, U.S.). Human lung adenocarcinoma cell line A549 were cultured in Dulbecco’s Modified Eagle Medium (DMEM, Gibco, U.S.) supplemented with 10% fetal bovine serum (FBS) and penicillin-streptomycin (ThermoFisher Scientific, Waltham, MA; 10 U.mL^–1^ and 100 µg.mL^–1^, respectively).

### Viruses

The HPAI H5N1 TX2/24 (A/Cattle/Texas/063224-24-1/2024, genotype B3.13, GISAID accession number: EPI_ISL_19155861) virus isolated from the milk of infected dairy cattle in Texas, USA(14) was used in animal studies. The TX2/24 virus was also used as a backbone to establish a reverse genetics system for generating recombinant TX2/24 (rTX2/24). The virus stock was propagated in 10-day-old embryonated chicken eggs (ECEs) and titrated in Cal-1 cells.

### Generation of recombinant viruses using reverse genetics technique

We used the reverse genetics technique to generate three recombinant HPAI H5N1 viruses belonging to clade 2.3.4.4b genotype B3.13, the bovine-derived rTX2/24 and two human-derived rTexas/37 and rMichigan/90. Initially, we used TX2/24 (A/Cattle/Texas/063224-24-1/2024, GISAID accession: EPI_ISL_19155861) as a backbone to establish the reverse genetic system and generate recombinant TX2/24 (rTX2/24). Briefly, full length genome sequences of the PB1, PB2, PA, HA, NA, NP, M and NS gene segments of the TX2/24 strain were chemically synthesized (Twist Bioscience) and cloned into the dual promoter influenza reverse genetics plasmid pHW2000 (kindly provided by Dr. Richard Webby at St. Jude Children’s Research Hospital) using the *BsmBI* (New England Biolabs) restriction sites. The pHW2000 plasmids containing eight gene segments (PB1, PB2, PA, HA, NA, NP, M and NS) of TX2/24 strain were co-transfected into a co-culture of HEK293T and Cal-1 (bovine uterine epithelial cells) using Lipofectamine 3000 reagents (ThermoFisher Scientific). Cell culture supernatant was harvested after 96 hours and used to infect newly seeded Cal-1 cells. Both cell lysate and culture supernatant were harvested after 72-96 hours to prepare the seed stock for the recombinant viruses. The working stock of the virus was prepared by inoculating 10-day-old embryonated chicken eggs (ECEs) via the allantoic cavity route and the infected allantoic fluid was harvested after 48 hours.

To generate the two human recombinant viruses (Texas/37 and Michigan/90), we aligned and compared the genome sequences of the human viruses (EPI_ISL_19027114 and EPI_ISL_19162802) with the bovine TX2/24 virus. Gene segments of the Texas/37 (PB2, PB1, PA and NS) and Michigan/90 (PB1, NA and NS) viruses that differ from the sequences of the TX2/24 virus were synthesized commercially (Twist Bioscience) and cloned into the pHW2000 vector as above. Internal *BsmBI* restriction sites (if any) were silenced by introducing synonymous mutations. Plasmid co-transfection and virus rescue (p0) in HEK293T-Cal-1 cells were performed as above. Preparation and sequencing of seed (p1) and working stocks (p2) were performed as above. Viruses from the initial rescue and from passages 1 and 2 were sequenced to confirm the identity of the respective wild-type virus sequences and the absence of unwanted mutations.

### Antiviral sensitivity analysis of the recombinant viruses *in vitro*

The antiviral sensitivity of wild-type TX2/24 and three recombinant viruses (rTX2/24, rTexas/37 and rMichigan/90) were evaluated against oseltamivir (neuraminidase inhibitor) and amantadine (M2 proton channel inhibitor) *in vitro*. MDCK cells seeded in 96-well plates were infected with each virus at a multiplicity of infection (MOI) of 0.1. Virus adsorption was carried out at 37 °C for 1 h, after which the inoculum was removed and replaced with 0.1 mL of DMEM supplemented with 2% fetal bovine serum containing oseltamivir or amantadine at concentrations ranging from 125 to 1000 nM. Plates were then incubated at 37 °C. Infected but untreated cells served as virus controls. At 24 h post-infection, culture supernatants were collected, and viral titers were determined in MDCK cells using endpoint dilution assays. Percent inhibition of viral titers relative to untreated controls was calculated to assess the relative sensitivity of each virus to the two antivirals.

### Replication kinetics of HPAI H5N1 viruses

Replication kinetics of HPAI H5N1 viruses were assessed in four mammalian cell lines: bovine mammary uterine epithelial cells (Cal-1), primary bovine mammary epithelial cells (bMEC), Madin-Darby Canine Kidney (MDCK) cells, and human lung adenocarcinoma cells (A549). Cells were seeded in 12-well plates at a density of 2.5x10^5^/mL and incubated at 37°C for 24 hours to reach approximately 90% confluency. Subsequently, cells were infected with one of the four H5N1 virus viruses - TX2/24, rTX2/24, rTexas/37, or rMichigan/90 - at a multiplicity of infection (MOI) of 0.1. Virus adsorption was performed at 4°C for 1 hour, after which the inoculum was removed and replaced with 1 mL of complete growth medium, and the plates were transferred to a 37°C incubator. Cells were incubated at 37°C and samples of cells plus supernatant were collected at 4, 8, 12, 24, 48, and 72 hours p.i., then stored at -80°C until use. Time point 0 was an aliquot of virus inoculum stored at -80°C as soon as inoculation was completed. Virus titers were determined in Cal-1 cells using endpoint dilution assays, with virus infectivity being determined by immunofluorescence assay (17) and titers were calculated using the Spearman and Karber’s method, expressed as TCID_50_.mL^–1^.

### Plaque morphology and phenotype

The plaque morphology and phenotype of H5N1 viruses were assessed in Cal-1, bMEC, MDCK, and A549 cells. Each cell type was seeded at a density of 5x10^5^ cells per well in 6-well plates and incubated for 24 hours at 37°C until they reached ∼90% confluency. Cells were then inoculated with one of four H5N1 viruses - TX2/24, rTX2/24, rTexas/37, or rMichigan/90 - at a target of 30 plaque-forming units (PFU) per well based on PFU titers determined in Cal-1 cells. Viral adsorption was carried out for 1 hour at 37°C. After incubation, the inoculum was removed, and each well was overlaid with 2 mL of a semi-solid medium containing 2X complete growth media mixed with 1% SeaKem agarose (resulting in a final concentration of 1X media and 0.5% agarose). Once the agarose had solidified, the plates were incubated at 37°C for 72 hours. After incubation, the agarose overlay was carefully removed. Cells were fixed with 3.7% formaldehyde for 30 minutes and stained with 0.5% crystal violet solution for 10 minutes at room temperature. Plaques were visualized and quantified using the ViralPlaque (37) software.

### Replicon assays

We performed replicon assays to evaluate the impact of amino acids variations in the influenza viral polymerase proteins on viral genome replication and gene transcription. We cloned viral polymerase proteins (PB2, PB1, PA) and NP of the bovine-derived TX2/24 and human-derived Texas/37 and Michigan/90 viruses into pcAGGS vector under the control of the chicken β-actin promoter. We chemically synthesized (GenScript, USA) two minigenome pUC57 plasmids encoding an influenza viral RNA-like segment expressing nanoluciferase (Nluc) or miniGFP2 flanked by the NP segment non-coding region (NCR). Of note, the NP segment of TX2/24, Texas/37 and Michigan/90 are identical. The transcription of the influenza vRNA is driven by a human RNA polymerase I (hPolI) promoter placed upstream of the NP(NCR)-NLuc or NP(NCR)-miniGFP cassettes. HEK293T cells were seeded in 24-well plate (1.25x10^5^/well) and co-transfected using PEI Prime (Sigma-Aldrich, USA) with 100 ng of each pcAGGS plasmids encoding the viral polymerase proteins (PB2, PB1, PA) and NP, and with 100 ng of the pUC57-NP(NCR)-Nluc or pUC57-NP(NCR)-miniGFP2 minigenome plasmid. Cells transfected with all influenza RNP proteins except the PB1-encoding plasmid served as a negative control. For luciferase assays, cell culture supernatant was removed after 24 and 48 hours and cells were lysed with 50 µL of 1X Passive Lysis Buffer (Promega) and Nluc expression was quantified using a Nano-Glo luciferase assay system (Promega, USA). For miniGFP2 expression, transfected cells were imaged daily using a fluorescence microscope (Olympus CKX53LED inverted microscope). The replicon assays were repeated four times independently.

### Pathogenesis studies in mice

C57BL/6 mice (*n* = 30) of 8 weeks of age were purchased from The Jackson Laboratory (United States) and housed in HEPA-filter isolators (Tecniplast BCU) with individual ventilation systems in the animal biosafety level 3 (ABSL-3) facility at the East Campus Research Facility (ECRF) at Cornell University. On day 0, mice (*n* = 6, 3 males, 3 females) were inoculated intranasally (under anesthesia) with 50 µL of virus suspension containing 1x10^3^ PFU of one of the following four viruses: TX2/24, rTX2/24, rTexas/37, or rMichigan/90. Mock-infected mice (*n =* 6) inoculated with cell culture media were kept as controls. Body weight, clinical signs, morbidity and mortality were recorded daily. Oropharyngeal swabs were collected using sterile rayon tip swabs on days 0, 1, 3 and 5 post-infection (pi) and placed in 1 mL virus transport medium (VTM Corning®, Glendale, AZ, USA) and stored at -80°C until processed for further analyses. Animals that reached humane endpoint (20% body weight loss) were euthanized and subjected to necropsy and tissue harvest. Tissues (brain, trachea, lungs, liver, spleen, small and large intestine) for virus quantification were collected aseptically in sterile containers and stored at -80°C until processed for further analyses. A section of each tissue was also collected in 10% neutral buffered formalin, processed for histopathological examination and stained routinely with hematoxylin and eosin (H&E). The study procedures were reviewed and approved by the Institutional Animal Care and Use Committee at Cornell University (IACUC approval number 2024-0094).

### Pathogenesis studies in hamsters

Pathogenesis evaluation of H5N1 viruses in hamsters was conducted in two phases. In the first phase, the wild type TX2/24 was compared with recombinant rTX2/24. A total of 21 7-8-week-old LVG golden Syrian hamsters (strain 049) were purchased from Charles River (United States) and housed in HEPA-filter isolators (Tecniplast BCU) with individual ventilation system in the ABSL-3 facility at ECRF at Cornell University. On day 0, hamsters were inoculated intranasally (under anesthesia) with 100 µL of virus suspension containing 5x10^5^ PFU of TX2/24 or rTX2/24 virus (*n* = 9/virus, 5 males and 4 females) and were housed individually in the BCU isolators. Mock-infected hamsters (*n =* 3) inoculated with cell culture media were kept as controls. Animals were monitored daily for clinical signs and body weight for 7 days. Oropharyngeal swabs were collected on days 0, 1, 3, 5 and 7 post-inoculation (pi). Upon collection, swabs were placed in sterile tubes containing 1 mL of viral transport medium (VTM Corning®, Glendale, AZ, USA) and stored at -80°C until processed for further analyses. Blood samples were collected on days 0 and 14 pi and sera were separated. Three animals from each inoculated group were euthanized under anesthesia on days 3, 5 and 7 pi and necropsied. Tissues (brain, olfactory bulb, nasal turbinate, trachea, lungs, liver, spleen, kidney, stomach, pancreas, intestine, uterus, ovary, testes, mandibular lymph node and tonsils) were collected in sterile tubes or placed in 10% neutral buffered formalin for virus isolation and histopathological analysis, respectively. A similar experimental design and sample collection timeline was followed while studying the pathogenesis of the two human H5N1 viruses, rTexas/37 and rMichigan/90. The study procedures were reviewed and approved by the Institutional Animal Care and Use Committee at Cornell University (IACUC approval number 2024-0094).

### Transmission studies in hamsters

The transmission studies in hamsters were conducted in two phases: TX2/24 virus was compared with rTX2/24 virus (phase I) and rTexas/37 was compared with rMichigan/90 virus (phase II). In each phase, 16 7-8-week-old LVG golden Syrian hamsters (strain 049) were purchased from Charles River (United States) and housed in HEPA-filter isolators as above. On day 0, hamsters were inoculated intranasally (under anesthesia) with 100 µL of virus suspension carrying 5x10^5^ PFU of TX2/24 or rTX2/24 virus (phase I) or rTexas/37 or rMichigan/90 (phase II) (*n* = 4/virus, 2 males and 2 females) and were housed individually in the BCU isolators. On the next day, each of the infected hamsters was transferred to a HEPA-filter isolator containing the contact animal and co-housed (1:1) for 48 hours. Animals were monitored daily for clinical signs and body weight for 14 days. Oropharyngeal swabs were collected daily from day 0-8 and on day 10 and 14 post-inoculation (pi) or post-contact (pc). Upon collection, swabs were placed in sterile tubes containing 1 mL of viral transport medium (VTM Corning®, Glendale, AZ, USA) and stored at -80°C until processed for further analyses. Blood samples were collected on days 0, 7 and 14 pi and sera were separated. Animals that died during the course of the experiment were necropsied on the day of death, while all surviving animals (infected and contact) were humanely euthanized and necropsied on day 14 pi. The study procedures were reviewed and approved by the Institutional Animal Care and Use Committee at Cornell University (IACUC approval number 2024-0094).

### RNA extraction and RT-PCR

Viral RNA from swab samples and tissue homogenates (10%, w/v) was extracted using the IndiMag Pathogen kit (INDICAL Bioscience) on the IndiMag 48s automated nucleic acid extractor (INDICAL Bioscience, Leipzig, Germany) following the manufacturer’s instructions. Real-time reverse transcriptase PCR (rRT-PCR) was performed using the Path-ID™ Multiplex One-Step RT-PCR Kit (Thermo Fisher, Waltham, MA, USA) and primers and probes targeting the M gene under the following conditions: 15 min at 48°C, 10 min at 95 °C, 40 cycles of 15 s at 95 °C for denaturation and 60 s at 60 °C. A standard curve was prepared using RNA extracted from HPAI H5N1 TX2/24 spiked media. Serial 10-fold dilutions of the stock virus (2x10^7^ log TCID_50_/mL) were prepared in DMEM, subjected to RNA extraction followed by RT-PCR as described above. The Ct values were used to estimate the viral RNA copy number in the tested samples using the relative quantification method.

### Virus isolation and titration

All oropharyngeal swabs and tissue homogenates were subjected to virus isolation under Biosafety Level 3 (BSL-3) conditions at the Animal Health Diagnostic Center (ADHC) Research Suite at Cornell University. Virus isolation was performed in bovine uterine epithelial cells (Cal-1). For virus titration, serial 10-fold dilutions of samples were prepared in MEM and inoculated into Cal-1 cells in 96-well plates. Two days later, culture supernatant was aspirated, and cells were fixed with 3.7% formaldehyde solution and subjected to IFA using the anti-NP (HB65) mouse monoclonal antibody, followed by secondary anti-mouse Alexa-594 incubation. The limit of detection (LOD) for infectious virus titration is 10^1.05^ TCID_50_.mL^–1^. Virus titers were determined at each time point using end-point dilutions and the Spearman and Karber’s method and expressed as TCID_50_.mL^–1^.

### Virus neutralization assay

Neutralizing antibodies in serum against HPAI H5N1 were assessed by virus neutralization (VN) assay using a recombinant TX2/24 virus expressing miniGFP2 (rTX2/24-miniGFP2) as previously described (17). For this, serial two-fold serum dilutions (1:8 to 1:1,024) of each serum sample were prepared in MEM and incubated with 200 TCID_50_ of rTX2/24-miniGFP2 for 1 h at 37 °C. Following that, 100 µL of Cal-1 cell suspension was added to each well of a 96-well plate and incubated at 37°C for 48 h. Plates were visualized using a fluorescence microscope (Hybrid microscope ECHO Revolve 3K) to determine neutralizing antibody (NA) titers, expressed as the reciprocal of the highest serum dilution capable of completely inhibiting HPAI H5N1 virus replication based on the expression of miniGFP2 by the rTX2/24-miniGFP2 virus.

### Histopathology

For the histological examination, tissue sections of approximately 0.5 cm^3^ were fixed by immersion in 10% neutral buffered formalin (≥20 volumes fixative to 1 volume tissue) for approximately 72 hours, and then transferred to 80% ethanol. These formalin-fixed paraffin-embedded tissues were processed, paraffin-embedded and sectioned at 5 µm thickness, stained with hematoxylin and eosin, and examined by a board-certified anatomic pathologist blinded to the experimental details. For all tissue, the degree of inflammation, necrosis, and hemorrhage was scored on the entirety of the section area as follows 0: none, 1: mild (affecting <25% of the tissue), 2: moderate (affecting >25% less than 50% of the tissue), 3: severe (affecting >% of the tissue). Additional tissue specific changes were scored similarly; brain: gliosis, satellitosis, neuronophagia; nasal turbinates, trachea, stomach small and large intestine, uterus: erosion/ulceration, granulation tissue; lung: pneumocyte type II hyperplasia, granulation tissue, fibrin; liver: granulation tissue, Kupffer cell hyperplasia and hypertrophy, granulation tissue; spleen: lymphoid hyperplasia, lymphoid depletion, congestion, extramedullary hematopoiesis; lymph node: lymphoid hyperplasia, lymphoid depletion, medullary cellularity.

### *In situ* RNA detection

Paraffin-embedded tissues were sectioned at 5 µm and subjected to *in situ* hybridization (ISH) using the RNAscope® ZZ probe technology (Advanced Cell Diagnostics, Newark, CA). Tissues from inoculated and controls were subjected to ISH using the RNAscope^®^ 2.5 HD Reagents–RED kit (Advanced Cell Diagnostics) and the V-InfluenzaA-H5N8-M2M1 probe (Advanced Cell Diagnostics), which targets H5Nx clade 2.3.4.4b viruses, as per the manufacturer’s instructions. ISH signals were amplified with multiple amplifiers conjugated with alkaline phosphatase enzymes, incubated with red substrate at room temperature for 10 min, and counterstained with hematoxylin.

### Statistical analysis and data plotting

Statistical analysis was performed by one-way or two-way analysis of variance (ANOVA) followed by multiple comparisons. Statistical analysis and data plotting were performed using the GraphPad Prism software (version 9.0.1).

## Acknowledgements

The work was supported by College of Veterinary Medicine Research Office and the Office of the Vice Provost for Research. DWB, RAA, and HCA were supported by NIH/NIAID grant 109022 to HCA.

## Supplementary Figures

**Supplementary Fig. 1.**
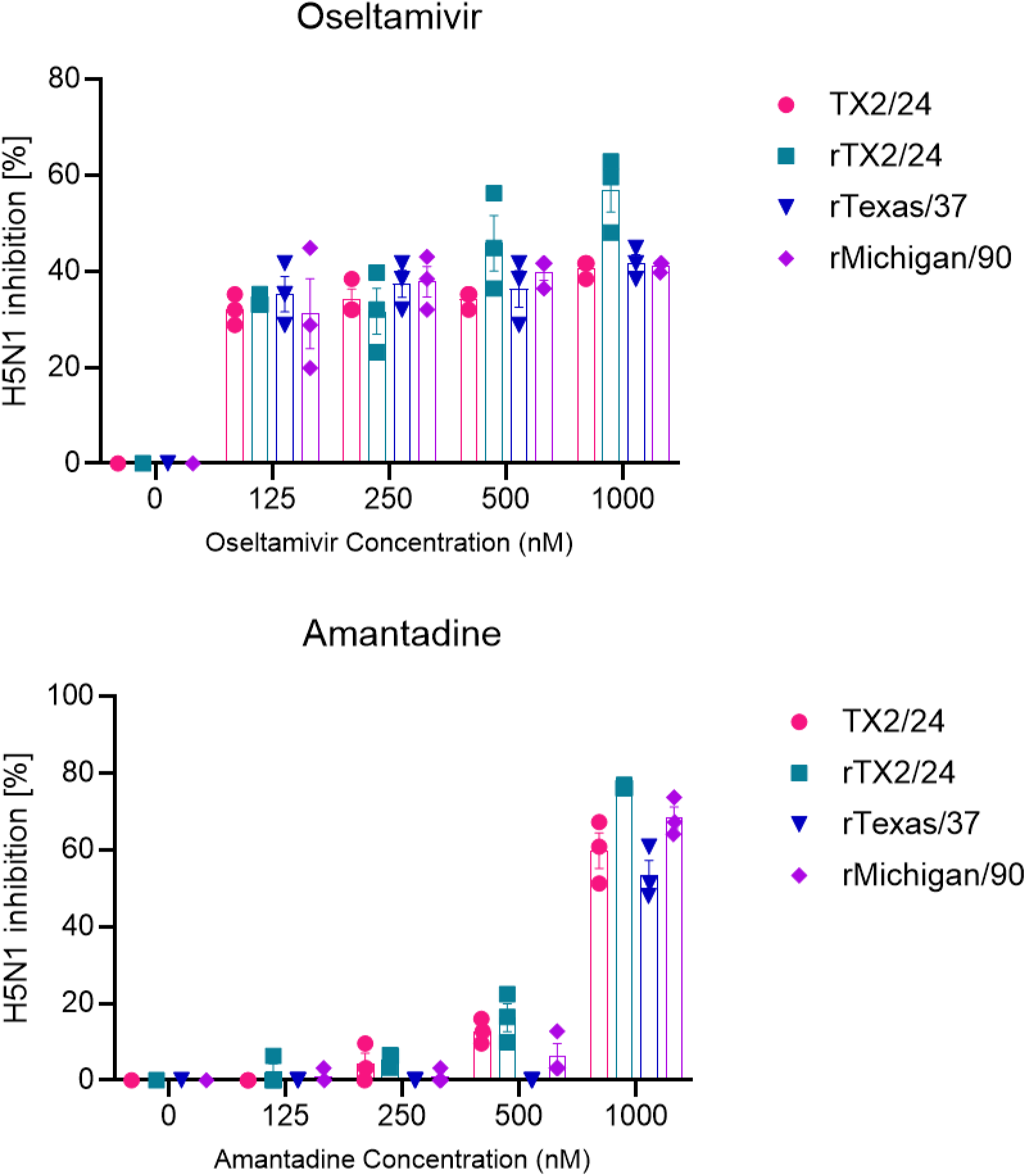
Antiviral sensitivity of H5N1 clade 2.3.4.4b viruses *in vitro*. MDCK cells seeded in 96-well plates were infected with either TX2/24, rTX2/24, rTexas/37 or rMichigan/90 virus at a multiplicity of infection (MOI) of 0.1. After 1 h virus adsorption at 37 °C, inoculum was removed and replaced with 0.1 mL of DMEM supplemented with 2% FBS containing oseltamivir or amantadine at concentrations ranging from 125 to 1000 nM. Culture supernatants were harvested after 24 h and virus titers were determined using endpoint dilutions and expressed as TCID_50_.mL^-1^. The limit of detection (LOD) for infectious virus titration was 10^1.05^ TCID_50_.mL^-1^. Data indicate mean ± SEM of 3 independent experiments.

**Supplementary Fig. 2.**
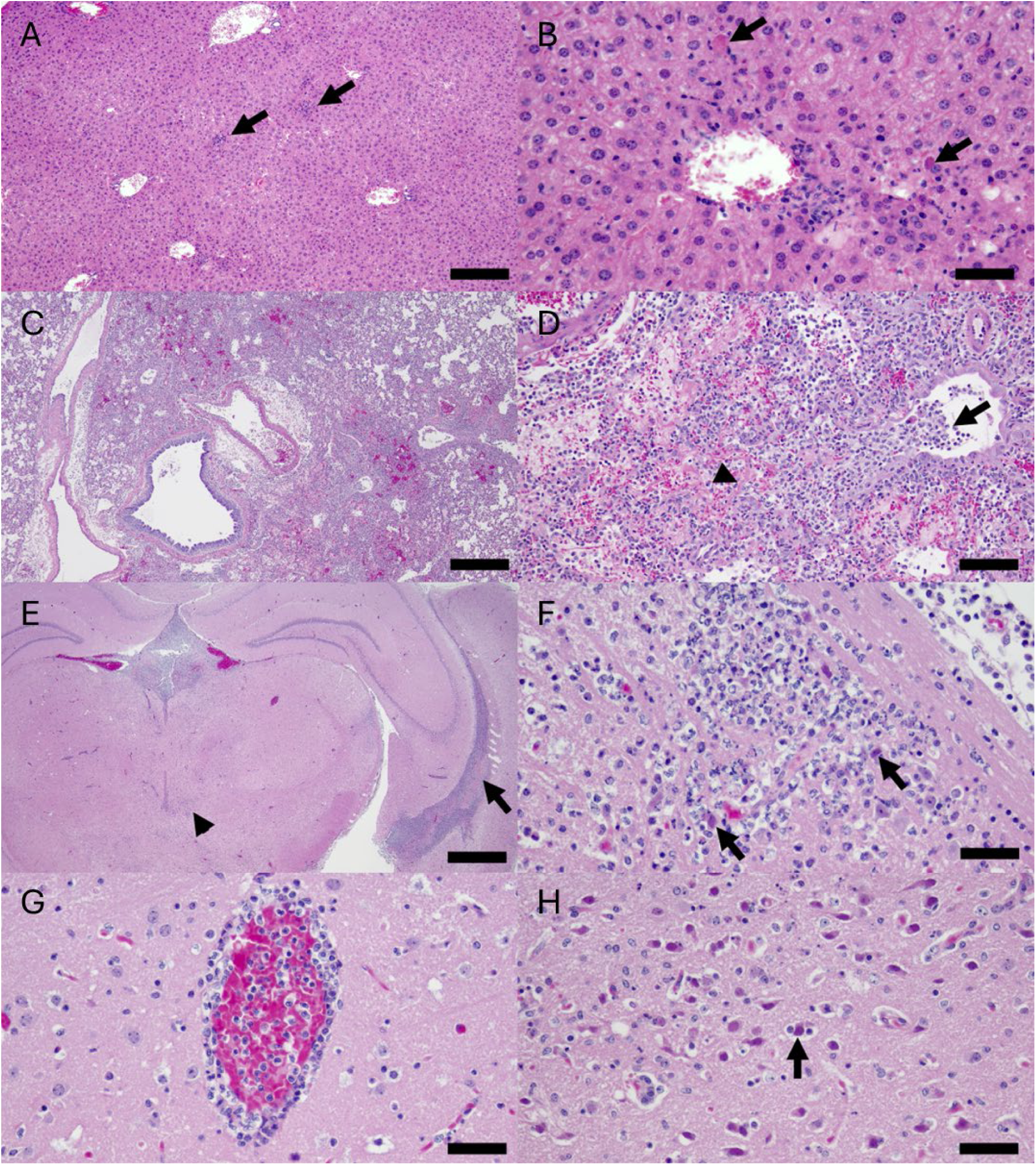
Histopathologic changes in mice and hamsters infected with influenza. A. Multifocal areas of inflammation and necrosis in a mouse liver. H&E, bar: 200 microns. B Higher magnification of an inflammatory focus from image A, highlighting neutrophils, macrophages, and lymphocytes along with necrotic hepatocytes (arrow). H&E, bar: 150 micros. C. Multifocal to coalescing areas of inflammation along with necrosis and hemorrhage in a hamster lung. H&E, bar: 200 microns. D. Mixed inflammation filling the lumen of a bronchiole (arrow) and expanding the interstitium, along with fibrin lining necrotic alveoli (arrowhead). H&E, bar: 100 microns. E. Mixed inflammation in the brain of a hamster filling a lateral ventricle (arrow) and in multiple random areas of the neuroparenchyma (arrowhead). H&E, bar: 250 micros. F. Higher magnification of an inflammatory focus in the neuroparenchyma from figure F, highlighting neutrophils, macrophages, lymphocytes, and glial cells along with necrotic neurons (arrows). G. Perivascular inflammatory cuff in the neuroparenchyma made up of lymphocytes and macrophages. H&E, bar: 100 microns. H. Necrotic neurons, occasionally flanked by lymphocytes (satellitosis, arrow). H&E, bar: 100 microns.

**Supplementary Fig. 3.**
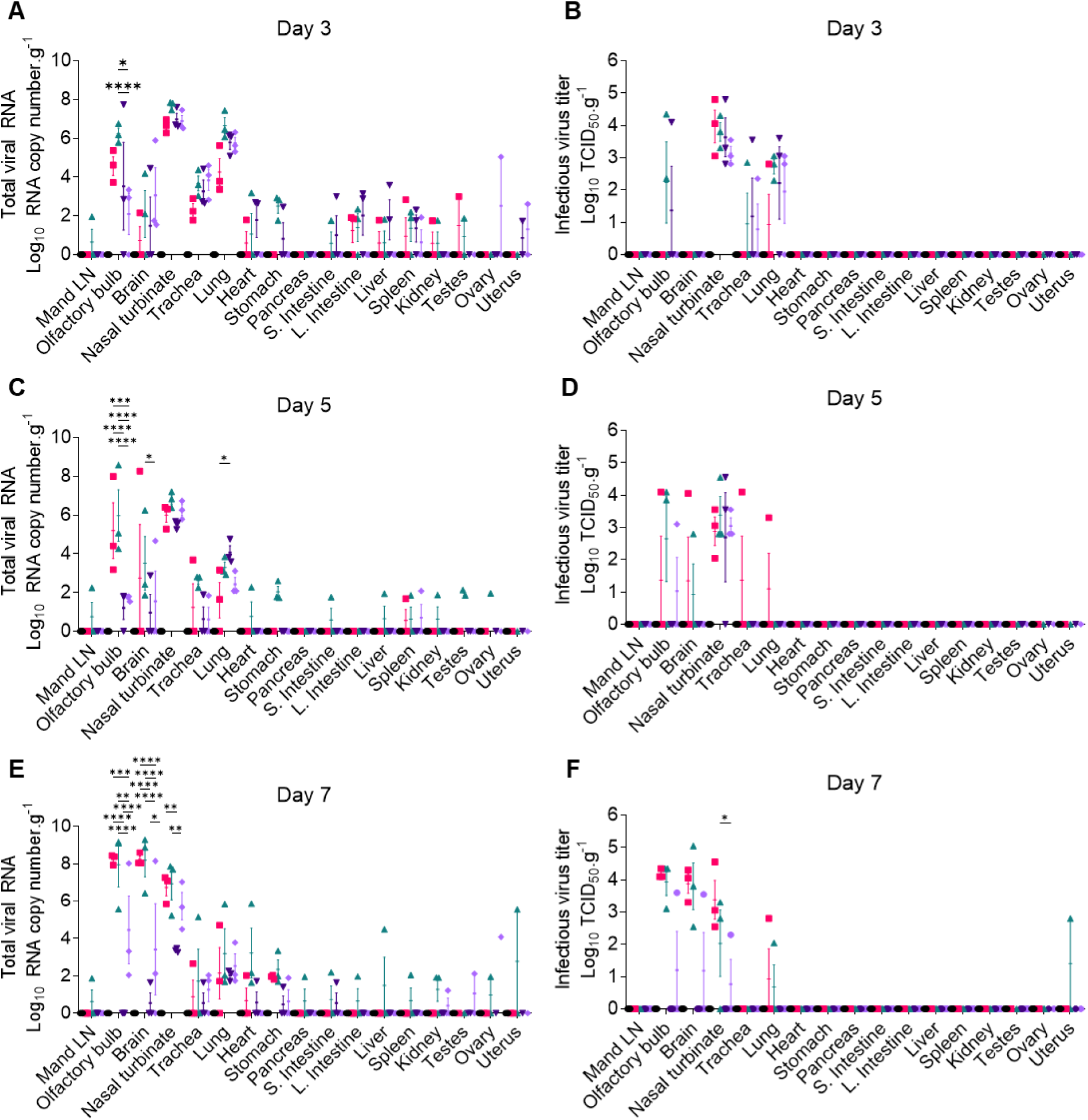
Virus distribution in tissues of hamsters infected with bovine- and human-derived HPAI H5N1 viruses. HPAI H5N1 viral RNA (A, C, E) and infectious virus (B, D, F) loads quantified by rRT-PCR and virus titration in tissues of hamsters collected at necropsy on days 3, 5 and 7 post-infection. Virus titers were determined using endpoint dilutions and expressed as TCID_50_.mL^-1^. The limit of detection (LOD) for infectious virus titration was 10^1.05^ TCID_50_.mL^-1^. Data indicate mean ± SEM of 3 mice per group per tissue per time point. Two-way ANOVA with Tukey’s multiple comparison test, * *p* < 0.05, ** *p* < 0.01, and **** *p* < 0.0001.

